# Subclonal Complete Loss of *CDKN1B* as a Common Genomic Alteration in Prostate Cancer: Associations with Race and Prostate Cancer Outcomes

**DOI:** 10.64898/2026.03.03.709424

**Authors:** Karen S. Sfanos, Rebecca Morton, Julia N. Flores, Rebecca Y. Sosa, Sarah E. Ernst, Luke Mummert, Jessica Hicks, Tamara L. Lotan, Jiayun Lu, Yuezhou Jing, Corrine E. Joshu, Angelo M. De Marzo

## Abstract

**Background:** Homozygous biallelic inactivation of *CDKN1B* is thought to be rare in cancer. Herein we evaluate the prevalence of intratumoral (subclonal) complete p27 protein loss (IPPL) in primary prostate cancer.

**Experimental Design:** We used immunohistochemistry (IHC) for p27 in a large cohort of whole tissue sections from radical prostatectomy (n=412) and metastases from self-identified African American (AA) and European American (EA) individuals. IPPL was evaluated alongside *CDKN1B* mRNA *in-situ* hybridization and next generation sequencing of laser captured cancer regions. Cox proportional hazards analyses assessed the association of IPPL with biochemical recurrence and development of metastases after radical prostatectomy.

**Results:** IPPL was detected in 18.1% of AA versus 12.2% of EA cases and was tightly correlated with *CDKN1B* mRNA loss and biallelic genomic loss. IPPL was associated with ≥pT3 pathologic stage and pN1 disease, however these associations were only significant among AA participants. IPPL was further associated in both univariate and multivariate analyses with the development of biochemical recurrence and metastasis after primary treatment, specifically in AA individuals. The prevalence of p27 genomic alterations in metastatic disease is higher than that of primary prostate cancer in publicly available datasets as well as our analysis of autopsy cases via IHC, indicating that complete p27 loss may be selected for in metastatic disease.

**Conclusions:** Subclonal biallelic loss of *CDKN1B* resulting in complete p27 protein loss is one of the most commonly occurring biallelic tumor suppressor genomic alterations in primary prostate cancer, and could contribute to worse prostate cancer outcomes, specifically in AA males.

## Introduction

Self-identified Black or African American (AA) males are more likely to be diagnosed with advanced prostate cancer and are nearly 2.5 times more likely to die from the disease than White/European American, Non-Hispanic (EA) males.^1^ The etiological factors driving this aggressive prostate cancer phenotype in AA males is likely multifactorial, and may include differences in access to care or social determinants of health as well as underlying genetics, diet and lifestyle factors, and environmental exposures.^2^ In addition, key somatic genomic driver alterations in prostate cancer occur at differing prevalence in AA and EA males, potentially contributing to disparities in clinical outcomes and suggesting that inclusion of diverse populations in genomic studies will improve precision medicine.^3–5^

Large-scale whole exome sequencing efforts, such as the TCGA, have uncovered the key somatic genomic drivers of prostate cancer, however these studies have largely been conducted with EA cohorts. Focused studies in AA cohorts have revealed markedly different frequencies of somatic alteration in several genes.^3–5^ For example, *ERG* rearrangements have been observed in approximately half of all prostate tumors occurring in EA males^6^, but these rearrangements are only seen in 25-30% of prostate cancers in AA males.^7,8^ The rate of some somatic alterations associated with poor outcomes are also lower in prostate cancer in AA males. For example, *PTEN* loss^3,8–10^ and *TP53* mutations^4,5,11^ are significantly lower in prostate cancer in AA compared to EA individuals. Conversely, *SPOP* mutations^4,12^, loss-of-function mutations in *ERF*^3^, mutations in *ZMYM3* and *FOXA1*^11^, and expression of *GSTP1*^13^ are more prevalent in prostate cancer in AA versus EA individuals. Finally, novel deletions in the *LSAMP* region were reported as enriched in AA patients, and were associated with biochemical recurrence (BCR).^14^ These data suggest that substantial differences in the prevalence of somatic genomic alterations may exist between prostate cancer in AA and EA individuals, however the key molecular drivers of worse outcomes in AA males remain to be elucidated.

*CDKN1B* (encoding p27) is a cyclin-dependent kinase (CDK) inhibitor that functions in controlling cellular division by binding to and preventing the activation of G_1_ cyclin-CDK complexes. In addition, p27 acts in numerous non-canonical, CDK-independent functions including cytoskeletal organization, cellular migration, and microtubule dynamics.^15^ *CDKN1B* is considered to be an atypical tumor suppressor that functions in a haplo-insufficient manner that does not generally require loss of function of both alleles according to the classical Knudson “two-hit” criteria. This was first described in p27 heterozygous mice that are predisposed to cancer at multiple sites when exposed to irradiation or a chemical carcinogen as well as the development of spontaneous pituitary cancer.^16^ The remaining wild-type allele was intact and not mutated in the cancer that developed in p27 heterozygous mice. Of keen interest, while loss of a single allele can drive increased cancer incidence, the incidence of cancers as well as mortality was increased in mice with homozygous *Cdkn1b* loss.^16^

The contribution of p27 to human cancer development and prognosis is somewhat enigmatic in accounting for its multiple forms of altered expression as well as its canonical and numerous non-canonical roles within the cell. Homozygous inactivating mutations and biallelic deletion of p27 in human cancer are considered rare.^17–19^ However, downregulation of p27 expression is found in many forms of cancer, and often associated with poor prognosis.^19,20^ Furthermore, mislocalization of the p27 protein from the nucleus to the cytoplasm, as is observed in some prostate cancers, is proto-oncogenic^19,21^, potentially via inhibition of RhoA activity, which in turn promotes carcinogenesis.^22^ In the normal prostatic epithelium, p27 protein is highly expressed in the nuclear compartment of luminal cells, but shows decreased nuclear expression in the putative precursor lesion high grade prostatic intraepithelial neoplasia as well as most invasive primary untreated adenocarcinomas.^23^ Genomic alterations of *CDKN1B* and decreased expression of p27 protein in prostate cancer have been extensively studied and there is a clear association between decreased expression or loss of p27 and aggressive prostate cancer. Lowered p27 expression (both nuclear and cytoplasmic) is associated with high tumor grade^23–26^ and biochemical recurrence (BCR).^25–30^ Furthermore, the combined loss of p27 and PTEN confers more aggressive disease in both animal models^31^ and human primary prostate cancer.^32^ The rate of *CDKN1B* alterations in metastatic disease is less clear, but homozygous deletion and/or loss of heterozygosity has been reported in as high as 40-44% of metastatic prostate cancer^33,34^ and reported to be homozygously or hemizygously deleted in circulating tumor DNA from 40 to over 50% of individuals with metastatic prostate cancer.^35,36^ This prevalence of p27 loss in metastatic disease is much higher than in localized cancer.^37–41^ Homozygous *CDKN1B* loss is thought to be rare in primary prostate cancer in both EA and AA indivduals. The prevalence of p27 alterations in primary prostate cancer in other races or ethnicities, such as Asian or Hispanic/Latino individuals, are understudied and unknown. Of keen interest, deep deletions of *CDKN1B* were previously found in 6.3% of 205 prostate tumors from AA individuals treated with radical prostatectomy, and were associated with an increased risk of metastasis in multivariate analysis.^37^ Herein, we report that the prevalence of subclonal *CDKN1B* genomic loss, especially among AA individuals, has been underestimated in primary prostate cancer. We further report that subclonal biallelic inactivation of *CDKN1B*, conferring complete p27 protein loss, in primary prostate cancer could contribute to worse prostate cancer outcomes, particularly in AA males.

## Materials and Methods

### Radical prostatectomy tissue specimens

All specimens were obtained and analyzed under an Institutional Review Board approved study. Formalin-fixed paraffin-embedded (FFPE) whole tissue slides (e.g. standard tissue slides) from tissue blocks were obtained from radical prostatectomy specimens (n = 412). The demographic and pathologic characteristics of the 412 cases are given in **Table 1**. The blocks chosen for this study contained the largest region of index tumor from the radical prostatectomy specimen. A single tissue block was assessed per case.

**Table 1.** Demographic and pathologic characteristics of radical prostatectomy whole tissue cohort.

|  | **AA** | **EA** | **Other** |
| --- | --- | --- | --- |
| **Race\*, # cases** | 210 | 197 | 5 |
| **Median Age, Years (Range)** | 59 (41-75) | 60 (40-74) | 67 (52-69) |
| **Gleason Grade Group, # cases** |  |  |  |
| 1 (%) | 57 (27.1) | 42 (21.3) | 0 |
| 2 (%) | 60 (28.6) | 68 (34.5) | 2 (40%) |
| 3 (%) | 29 (13.8) | 40 (20.3) | 2 (40%) |
| 4-5 (%) | 64 (30.5) | 47 (23.9) | 1 (20%) |
| **Stage, # cases** |  |  |  |
| Organ confined (pT2), N0 (%) | 116 (55.2) | 103 (52.6)** | 3 (60%) |
| ≥pT3 and/or N1 (%) | 94 (44.8) | 93 (47.4) | 2 (40%) |
| N0 (%) | 172 (81.9) | 162 (82.7) | 4 (80%) |
| N1 (%) | 38 (18.1) | 34 (17.3) | 1 (20%) |

\*Self-identified race
\*\*Stage could not be assigned to one EA participant

### Tissue microarray (TMA) sets

The “PCBN High Grade Race” TMA set was assembled from the same blocks as 180 of the cases used for whole tissue analyses. The PCBN High Grade Race TMA set was used to compare detection rates of intratumoral complete p27 protein loss in TMAs to the rate of loss within whole tissue sections. This TMA set contains radical prostatectomy tumor tissue from 120 self-identified AA males matched 2:1 to 60 EA males on age +/-3 years, grade and stage and contains 4 × 0.6mm tissue spots of the index tumor from each case.

The “Metastasis at Autopsy” TMA contains tissues from autopsy metastatic sites from 2 AA and 4 EA males. The details of the Metastasis at Autopsy TMA set have been previously described.^42^

### Immunohistochemistry (IHC) and IHC scoring

p27 IHC was performed on the Ventana Discovery Ultra IHC/ISH platform (Roche Diagnostics) using an antibody for p27 (BD Biosciences, clone 57/Kip1/p27). The p27 antibody has been extensively validated in our laboratory as previously described.^43^ IHC for TP53 (p53), PTEN, and ERG was performed as previously reported in a subset of the cases.^44,45^ IHC-stained slides were initially screened for p27 loss using a light microscope and then the slides were scanned using a Ventana DP 200 slide scanner (Roche Diagnostics). Slide images were housed in a Concentriq LS repository (Proscia) where they were screened for a second time for p27 loss. Determination of p27 loss required an internal positive control of neighboring cancer cells and/or non-cancer cells within the stroma of p27-cancer regions maintaining p27+ staining.

### RNA in situ hybridization (RISH)

Chromogenic RISH was performed using the RNAscope 2.5 FFPE Brown Reagent Kit (Cat. No. 322310, Advanced Cell Diagnostics, Newark, CA) per the manufacturer instructions. RNAscope target probes used were designed against *CDKN1B* (p27) mRNA (Cat. No. 607061) or peptidyl prolyl isomerase B (PPIB), also known as cyclophilin B, as a positive control mRNA (Cat. No. 313901).

### DNA and RNA extraction from FFPE tissues

FFPE tissues were cut onto 5 membrane slides for laser microdissection (LCM) with a Leica LMD 7000 instrument or macro-dissection with a scalpel, depending on the size of the p27 loss region. DNA/RNA were simultaneously extracted from micro- or macro-dissected tissues using the MagMAX™ FFPE DNA/RNA Ultra Kit (ThermoFisher Scientific). Extracted DNA was quantified using the Qubit dsDNA High Sensitivity kit (ThermoFisher).

### Next generation sequencing (NGS) panel sequencing and analysis

DNA samples from matched benign, regions of adenocarcinoma staining positive for p27 protein (p27+ cancer), and regions of adenocarcinomas showing complete loss of p27 protein (p27-cancer) from 7 cases were submitted to the to the Molecular Diagnostics Laboratory in the Department of Pathology at Johns Hopkins Hospital for analysis on the NGS solid tumor panel. Library preparation was performed using Kapa Roche reagents and hybrid capture was performed using Integrated DNA Technologies probes. Sequencing was performed to an average unique read depth of greater than 500X using Sequencing by Synthesis 2 × 100 base pair paired-end cluster generation on the Illumina NovaSeq 6000 platform. FASTQ files were generated from Binary Cluster Files (.bcl) using the Illumina bcl2fastq v1.8.4 software with parameters set as per manufacturer’s instructions. FASTQ files were aligned to the human genome reference hg19 (GRCh37) using the Burrows-Wheeler Aligner v0.7.10 algorithm with default settings. BAM files were generated using Picard Tools v1.119 and variant calling is performed using in-house variant caller algorithm (MDLVC v5.0) cross-referenced with HaplotypeCaller (Genome Analysis Tool Kit 3.3) under discovery mode in the coding regions of target genes. All variant calls are evaluated with Integrated Genomics Viewer v2.3.4 (IGV; Broad Institute, MIT Harvard, Cambridge, MA, USA) and annotated with dbSNP v150 and COSMIC v82 databases.

### cBioPortal analyses

cBioPortal^46^ data was used to assess mutation and deep deletion frequency in the *CDKN1B* gene in prior genomic studies of prostate cancer. The following studies were used to determine frequency assessments and analyses by race: Race Differences in Prostate Cancer (MSK, 2021), Prostate Cancer (MSK, Clin Cancer Res 2024), Prostate Adenocarcinoma (MSK, Clin Cancer Res 2022), and Prostate Adenocarcinoma (MSK, Eur Urol 2020). The following studies were used to assess reported mutations across all prostate cancer studies: Race Differences in Prostate Cancer (MSK, 2021), Prostate Adenocarcinoma (MSK, Clin Cancer Res. 2022), Prostate Cancer (MSK, Clin Cancer Res 2024), Prostate Cancer (MSK, Science 2022), Prostate Adenocarcinoma (TCGA, Firehose Legacy), Metastatic Prostate Adenocarcinoma (MCTP, Nature 2012), Prostate Adenocarcinoma (TCGA, Cell 2015), Prostate Cancer (MSK, JCO Precis Oncol 2017), Prostate Adenocarcinoma (TCGA, PanCancer Atlas), Metastatic castration-sensitive prostate cancer (MSK, Clin Cancer Res 2020), Prostate Adenocarcinoma (MSK, Eur Urol 2020), The Metastatic Prostate Cancer Project (Provisional, June 2021), Prostate Adenocarcinoma (CPC-GENE, Nature 2017), Metastatic Prostate Adenocarcinoma (SU2C/PCF Dream Team, PNAS 2019), Prostate Adenocarcinoma (TCGA, GDC), Prostate Adenocarcinoma (Broad/Cornell, Nat Genet 2012), Metastatic Prostate Cancer (SU2C/PCF Dream Team, Cell 2015), Prostate Adenocarcinoma (SMMU, Eur Urol 2017), Prostate Cancer MDA PCa PDX (MD Anderson, Clin Cancer Res 2024), and Prostate Adenocarcinoma (MSK/DFCI, Nature Genetics 2018).

### Statistical analyses

Comparison of p27 protein status by IHC by race, Gleason score, pathologic stage, PTEN loss, ERG positivity, and TP53 mutation status were performed by Chi-Square test or Fisher’s Exact test. Cox proportional hazards regression was used to estimate the hazard ratio and 95% CI of biochemical recurrence and development of metastases after radical prostatectomy by p27-loss status. Analyses were further adjusted for race (EA versus AA), grade (prostatectomy Gleason grade group 1,2,3, or 4-5), stage (pT2 or ≥pT3), and lymph node status (pN0 or pN1). To test whether the association between p27 and biochemical recurrence and/or metastases is modified by race, we (1) stratified analyses by race and (2) used the likelihood ratio test to test the statistical significance of an interaction term between p27 loss and race in the main models. All tests were two-sided, with *p* < 0.05 considered to be statistically significant. All were conducted using SAS 9.4 (SAS Institute, Cary, NC, USA).

### Data availability

The data generated in this study are available within the article and the supplementary data files.

## Results

### Identification of subclonal complete loss of CDKN1B in primary prostate cancer

We routinely use p27 IHC to assess the suitability of donor paraffin blocks when constructing TMAs, as p27 IHC is sensitive to how well a tissue has been fixed.^43^ During construction of a series of TMA sets inclusive of matched AA and EA individuals with primary prostate cancer, we noted that many of the cancers exhibited subregions of complete loss of staining for p27 (p27-) by IHC that were often present as foci within larger tumor regions that showed positve staining for p27 (p27+) in the remainder of the tumor cells (**Fig. 1A**). We will refer to these p27-foci as intratumoral complete p27 protein loss (IPPL) regions. The p27+ adenocarcinoma directly adjacent to the IPPL regions indicated that the lack of p27 staining was not simply an artifact related to poor tissue fixation. In addition, as a further internal positive control, in all regions that we classified as IPPL, non-cancer cells within the stroma of p27-cancer regions maintained p27+ staining (**Fig. S1**). Furthermore, the use of RISH for *CDKN1B* mRNA demonstrated that p27 protein loss observed via IHC was also reflective of loss at the mRNA level (**Fig. 1B**).

**Figure 1.**
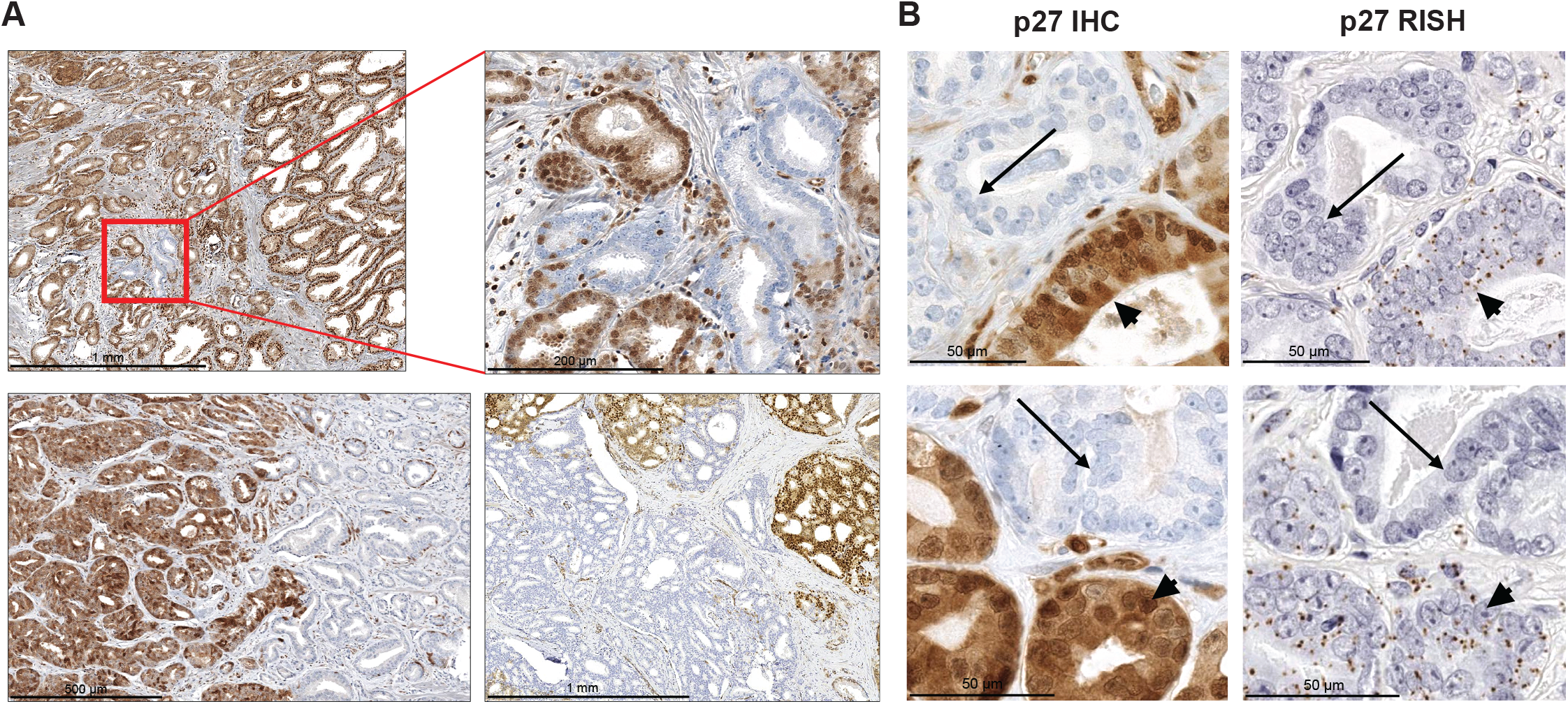
Examples of intratumoral complete loss of p27 protein in primary prostate cancer assessed via IHC and RISH. **A**) p27 IHC in primary prostate cancer. Brown staining is p27-positive cancer cells. Blue staining is nuclear hematoxylin counterstaining in cancer cells that are completely devoid of p27 protein. Top right panel is a higher magnification of the red boxed area in the top left panel. **B**) p27 IHC versus *CDKN1B* RISH on adjacent sections of primary prostate cancer. Note the areas negative for p27 IHC (arrows) are negative for *CDKN1B* mRNA by RISH (arrows) whereas areas positive for p27 by IHC (brown stain, arrowheads) are positive for *CDKN1B* mRNA by RISH (brown dots, arrowheads).

### Confirmation of genomic loss of p27

We dissected FFPE tissue sections for benign, p27+ cancer, and p27− cancer regions from seven cases where we identified IPPL. DNA was extracted from the dissected tissues for use with an NGS solid tumor panel that includes *CDKN1B* and can assess point mutations, small insertion/deletion mutations, and copy number alterations. The results of panel sequencing revealed that areas that were negative for p27 protein via IHC contained either somatic mutations in *CKDN1B* (presumably silencing mutations), genomic deletions, or a combination of the two (**Fig. 2A,B**). Of interest, in three cases, somatic genomic alterations in *CDKN1B* (that were absent in benign tissues) were also noted in the p27+ cancer regions. This finding indicates that, as expected, hemizygous alterations can occur somatically within *CDKN1B* without complete abrogation of protein expression in prostate cancer. Interestingly, we noted that some of these regions contained mis-localized (e.g., predominantly cytoplasmic) and/or lowered p27 protein expression (**Fig. S2**), however mis-localized and lowered p27 protein expression was also observed in some p27+ cancer regions without detected p27 genomic alterations (**Table S1**). This latter observation should be interpreted carefully, however, due to 1) limitations to the sensitivity of the panel sequencing based on tumor purity and 2) potential artifact from poor fixation^43^, although our internal positive control of non-cancer cells within the stroma with strong nuclear p27+ staining was maintained (**Fig. S2**). An analysis of reported mutations involving *CDKN1B* in cBioPortal^46^ among all prostate cancer studies revealed that mutations occur along the entire gene, with close to half of reported mutations occurring in the cell cycle inhibitory region (**Fig. 2C**). Consistent with this finding, 2/4 of the mutations identified in the current study occurred in the cell cycle inhibitory region. The majority of reported mutations are truncating mutations, also consistent with those identified in the present study (**Fig. 2C**).

**Figure 2.**
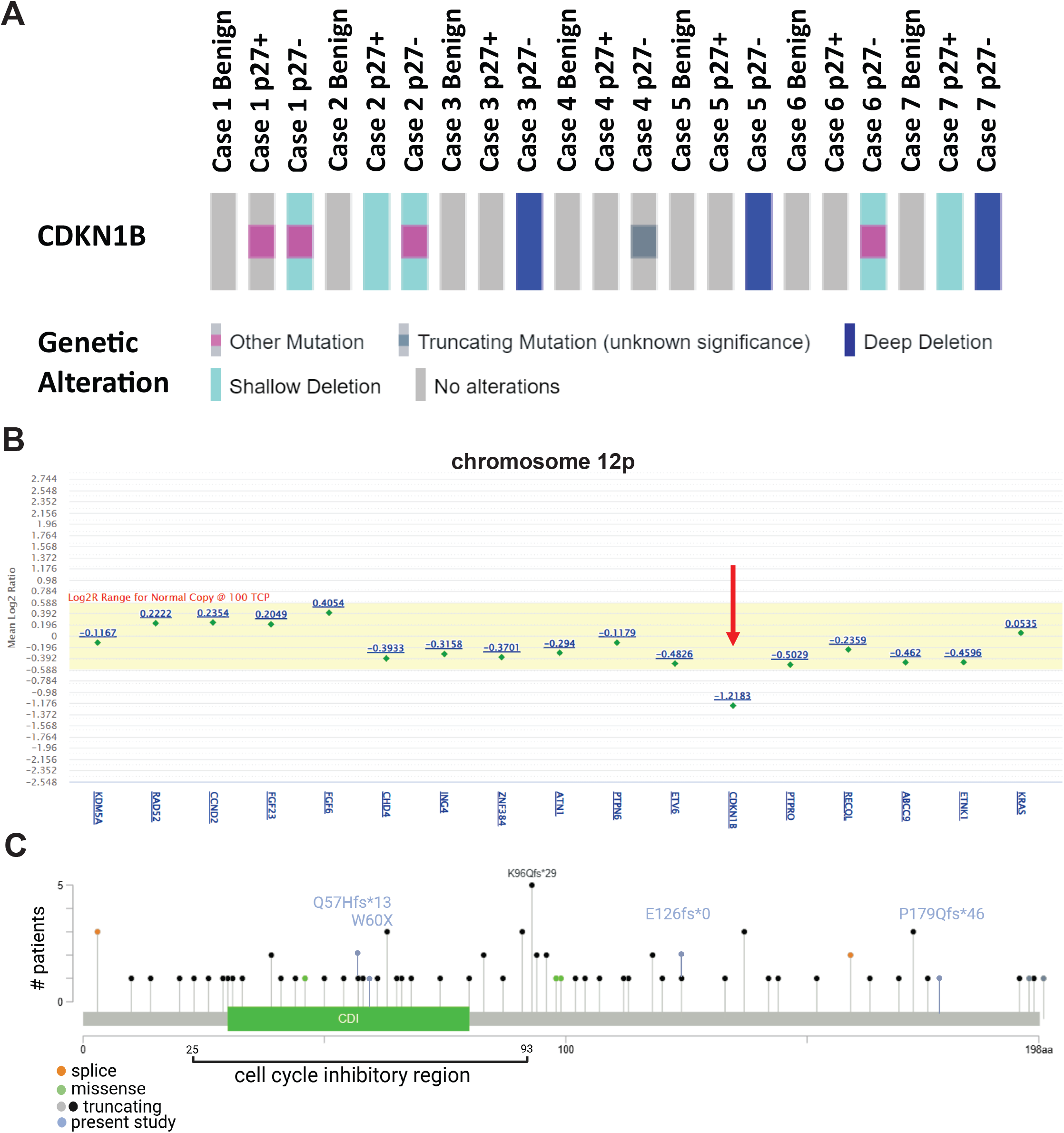
Patterns of genomic loss of *CDKN1B* in primary prostate cancer. **A**) *CDKN1B* genomic alterations in seven primary prostate cancer cases as assessed by NGS panel sequencing of individually dissected regions. p27+ = regions of adenocarcinoma staining positive for p27 protein, p27− = regions of adenocarcinoma staining negative for p27 protein. **B**) Example of copy number variant (CNV) analysis of chromosome 12p using in-house CNV caller software (MDL VC v10.0) showing deletion of the *CDKN1B* gene (p27, red arrow) in case 5. **C**) Lollipop plot of reported mutations across all prostate cancer studies in cBioPortal. Mutations identified in the present study (n=4) are indicated in light blue. CDI=cyclin-dependent kinase inhibitor.

### Prevalence of IPPL by race and pathologic factors

Inactivating mutations and deep deletions of the *CDKN1B* gene have been previously reported in prostate cancer, but at a low frequency. A prior study of 205 primary prostate cancers in AA individuals reported deep deletions in *CDKN1B* in 6.3% of the cases.^37^ Likewise, the frequency of *CDKN1B* genomic alterations (deep deletions, mutations, splice variants) reported in primary prostate cancer in datasets available in cBioPortal^46^ range from ~2-4% of cases (**Table S2**). When separated by race (AA vs EA), *CDKN1B* alterations were consistently higher in primary prostate cancer in AA (4.8-7.9%) versus EA cases (2.9-4.0%), although not statistically significant (**Table S3**). Due to the subclonal nature of IPPL that we observed in the cases in the present study that sometimes only encompassed small regions comprised of a few cancer glands (**Fig. 1A**), we reasoned that this alteration would almost certainly often go undetected in sequencing strategies that assess bulk tumor tissues. In other words, unless the IPPL region was expansive or microdissected from the tissue, as we did in our NGS panel sequencing studies (**Fig. 2**), the alterations in *CDKN1B* would be missed during NGS analyses since the majority of reads would presumably come from p27 intact tumor cells (or from cancer cells with single allele alterations only) and normal stromal/other cells.

Due to this potential underestimation of *CDKN1B* genomic alterations in published datasets, we expanded our cohort to a total of 412 whole tissue sections (self-identified 210 AA, 197 EA, 5 other race) that we assessed for p27 via IHC (**Table 1**). Careful examination of these tissue sections revealed a prevalence of IPPL of 15.3% among all cases (**Table 2**). The prevalence of IPPL was higher in primary prostate cancer from AA cases (18.1%) versus EA cases (12.2%), although this did not reach statistical significance (**Table 2**). Of the cases where IPPL was identified, 61.3% were from AA individuals versus 38.7% from EA individuals. This rate of IPPL in primary prostate cancer is 3-5 times higher than the previously reported prevalence of deep deletions in radical prostatectomy specimens from AA individuals^37^ or in datasets contained in cBioPortal (**Tables S2-S3**). In further support of the concept that the frequency of IPPL may be underestimated due to its sublconal nature, whereas we detected IPPL in 15.8% (28/177) of whole tissue sections assessed via p27 IHC, we only detected IPPL in 2.9% (5/175) of the same cases sub-sampled for a TMA set and assessed via p27 IHC (**Fig. S3**).

**Table 2.**
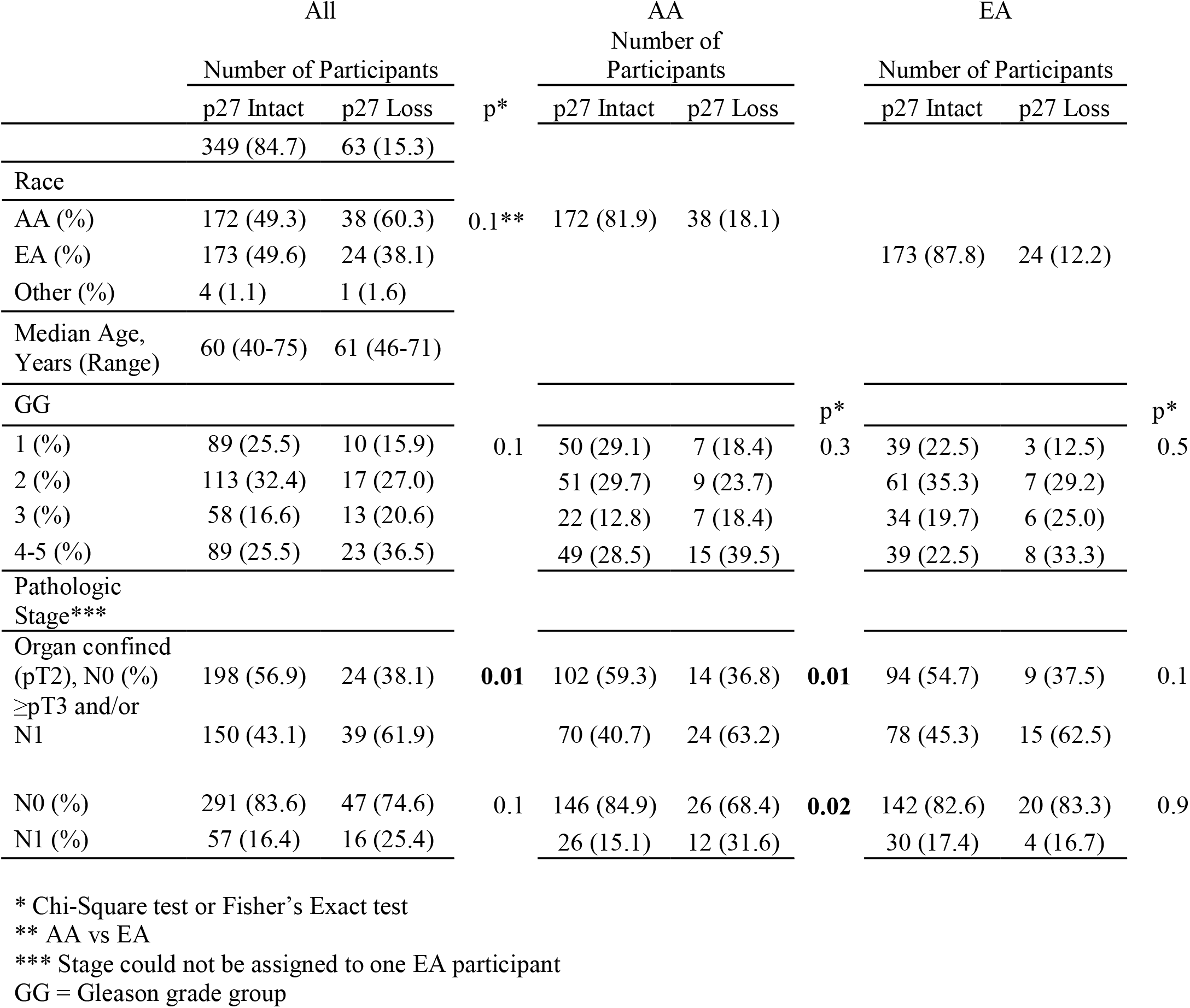
Prevalence of p27 loss (IPPL) by race and pathologic factors.

Among all cases, IPPL was not significantly associated with higher Gleason grade group (p=0.1) or pathologic lymph node involvement at time of prostatectomy (pN1 versus pN0, p=0.1) but was significantly associated with pathologic stage when assessed as ≥pT3 disease and/or pN1 versus organ confined (pT2), N0 disease (p=0.01, **Table 2**). Interestingly, when cases were examined separately by race (AA vs EA), a significant association with stage (p =0.01) and pN1 disease (p=0.02) was only found among AA participants (**Table 2**).

### IPPL is not correlated with other common genomic alterations in prostate cancer

A subset of the cases included in the present study had previously been screened for PTEN loss, ERG positivity, and TP53 mutation via IHC.^44,45^ An analysis of these common genomic alterations in primary prostate cancer in association with the detection of IPPL revealed no statistically significant associations (**Table S5**).

### IPPL is correlated with adverse outcomes in AA males

Outcomes data for the development of biochemical (PSA) recurrence and metastasis after radical prostatectomy was available for a subset of the study participants (n=222). There was a positive association with biochemical recurrence when all participants were assessed in a Cox hazard regression model without any adjustment (p=0.05, **Table 3**), however adjustment for race (EA vs AA), age, stage (organ confined (pT2), N0 vs ≥pT3 and/or N1) and Gleason grade group (1, 2, 3, 4-5) attenuated the association. When the Cox hazard regression model was separately analyzed by race, there was no association between IPPL and biochemical recurrence among EA participants, however there was a strong association between IPPL and increased risk of biochemical recurrence among AA participants (HR 5.59, 95% CI 2.32-13.48, p=0.0001) that was also significant when adjusted for age, stage (organ confined (pT2), N0 vs ≥pT3 and/or N1) and Gleason grade group (HR 3.05, 95% CI 1.09-8.51, p=0.03) as well as adjustment for age, stage (N0 vs N1) and Gleason grade group (HR 3.71, 95% CI 1.19-11.56, p=0.02, **Table 3**). Likewise, the interaction term for IPPL and race in the fully adjusted model was statistically significant (p-for interaction =0.04).

**Table 3.** Hazard ratio (HR) of p27 status on prostate cancer biochemical recurrence.

| All | Case<br>(N) | Control<br>(N) | HR (95% CI) * |  |  | *p* | HR (95% CI) ** |  |  | *p* | HR (95% CI) *** |  |  | *p* |
| --- | --- | --- | --- | --- | --- | --- | --- | --- | --- | --- | --- | --- | --- | --- |
| P27 Intact | 49 | 142□ | Ref |  |  | 0.05 | Ref |  |  | 0.8 | Ref |  |  | 0.6 |
| P27 Loss | 13 | 17 | 1.85 | 1 | 3.41 |  | 1.08 | 0.57 | 2.08 |  | 1.19 | 0.61 | 2.32 |  |
| EA |  |  |  |  |  |  |  |  |  |  |  |  |  |  |
| P27 Intact | 37 | 68□ | Ref |  |  | 0.5 | Ref |  |  | 0.3 | Ref |  |  | 0.5 |
| P27 Loss | 3 | 11 | 0.66 | 0.2 | 2.15 |  | 0.51 | 0.15 | 1.76 |  | 0.66 | 0.2 | 2.18 |  |
| AA |  |  |  |  |  |  |  |  |  |  |  |  |  |  |
| P27 Intact | 10 | 73 | Ref |  |  | **0.0001** | Ref |  |  | **0.03** | Ref |  |  | **0.02** |
| P27 Loss | 10 | 6 | 5.59 | 2.32 | 13.5 |  | 3.05 | 1.09 | 8.51 |  | 3.71 | 1.19 | 11.6 |  |

\* HR and 95% CI were estimated from Cox hazard regression model without any adjustment.
\*\*, HR and 95% CI were estimated from Cox hazard regression model adjusting for age, stage (organ confined (pT2), N0 vs ≥pT3 and/or N1), Gleason grade group (1,2,3,4-5), and race (if applicable).
\*\*\*, HR and 95% CI were estimated from Cox hazard regression model adjusting for age, stage (N0 vs N1), Gleason grade group (1,2,3,4-5), and race (if applicable).
□ One EA participant without assigned stage was excluded from adjusted models

There was a significant association between IPPL and the development of metastasis when all participants were assessed in a Cox hazard regression model without any adjustment (HR 3.15, 95% CI 1.22-8.14, p=0.02); this association remained positive but was no longer statistically significant after adjustment for race and clinical factors (**Table 4**). When stratified by race, again there was no significant association between IPPL and metastasis among EA participants, but there was an association between IPPL and increased risk of metastasis among AA participants (HR 19.01, 95% CI 2.11-171.24, p=0.01, **Table 4**); this association was positive but not statistically significant upon further adjustments for age, stage, and grade. Similarly, the interaction term for IPPL and race in a race-adjusted model was statistically significant (p-for interaction =0.04) but attenuated in the fully adjusted model (p-for interaction =0.1). Of note, the number of metastatic events were limited in the current data set.

**Table 4.** Hazard ratios (HR) of p27 status on prostate cancer metastasis.

| All | Case<br>(N) | Control(N) | HR (95% CI) * |  |  | *p* | HR (95% CI) ** |  |  | *p* | HR (95% CI) *** |  |  | *p* |
| --- | --- | --- | --- | --- | --- | --- | --- | --- | --- | --- | --- | --- | --- | --- |
| P27 Intact | 15 | 177□ | Ref |  |  | **0.02** | Ref |  |  | 0.2 | Ref |  |  | 0.1 |
| P27 Loss | 6 | 24 | 3.15 | 1.22 | 8.14 |  | 2.04 | 0.72 | 5.81 |  | 2.63 | 0.89 | 7.78 |  |
| EA |  |  |  |  |  |  |  |  |  |  |  |  |  |  |
| P27 Intact | 14 | 91□ | Ref |  |  | 0.7 | Ref |  |  | 0.9 | Ref |  |  | 0.5 |
| P27 Loss | 2 | 12 | 1.38 | 0.31 | 6.13 |  | 1.16 | 0.23 | 5.85 |  | 1.65 | 0.35 | 7.72 |  |
| AA |  |  |  |  |  |  |  |  |  |  |  |  |  |  |
| P27 Intact | 1 | 84 | Ref |  |  | **0.01** | Ref |  |  | 0.1 | Ref |  |  | 0.3 |
| P27 Loss | 4 | 12 | 19.01 | 2.11 | 171 |  | 7.17 | 0.54 | 95.4 |  | 3.46 | 0.34 | 35.4 |  |

\* HR and 95% CI were estimated from Cox hazard regression model without any adjustment.
\*\*, HR and 95% CI were estimated from Cox hazard regression model adjusting for age, stage (organ confined (pT2), N0 vs ≥pT3 and/or N1), Gleason grade group (1,2,3,4-5), and race (if applicable).
\*\*\*, HR and 95% CI were estimated from Cox hazard regression model adjusting for age, stage (N0 vs N1), Gleason grade group (1,2,3,4-5), and race (if applicable).
□ One EA participant without assigned stage was excluded from adjusted models

### p27 protein loss in metastatic prostate cancer

Due to our finding that IPPL is associated with higher stage prostate cancer, as well as worse outcomes including the development of metastases after primary treatment with radical prostatectomy, we next questioned whether p27 protein loss is prevalent among metastatic prostate cancer. Prior studies have detected homozygous deletion and/or loss of heterozygosity in tissue^33,34^ and circulating tumor DNA^35,36^ from close to half of metastatic prostate cancer in predominantly EA cohorts. In an analysis of data available in cBioPortal, we found that the detection of mutations/structural variants as well as homozygous deletions in *CDKN1B* was significantly higher in metastatic samples versus primary prostate cancer (**Table S2**), with over twice as many alterations found in metastatic samples. Furthermore, in a dataset designed to assess race differences in prostate cancer, we noted a significantly higher prevalence of deep deletions involving the *CDKN1B* gene in AA versus EA participants (**Table S3**). We performed p27 IHC on a TMA set comprised metastatic sites collected at autopsy from 2 AA and 4 EA individuals. p27 loss was detected in 4 of the 6 autopsy cases (**Table S4**) and p27 loss was detected in metastases from both AA individuals represented on the TMA. Interestingly, in most cases, p27 loss was only detected in a subset of metastases. Unlike primary prostate cancer, however, we observed cases with p27 loss across all cancer cells in a metastatic site (**Fig. 3**), suggesting clonal selection for cancer cells with p27 protein loss at that site.

**Figure 3.**
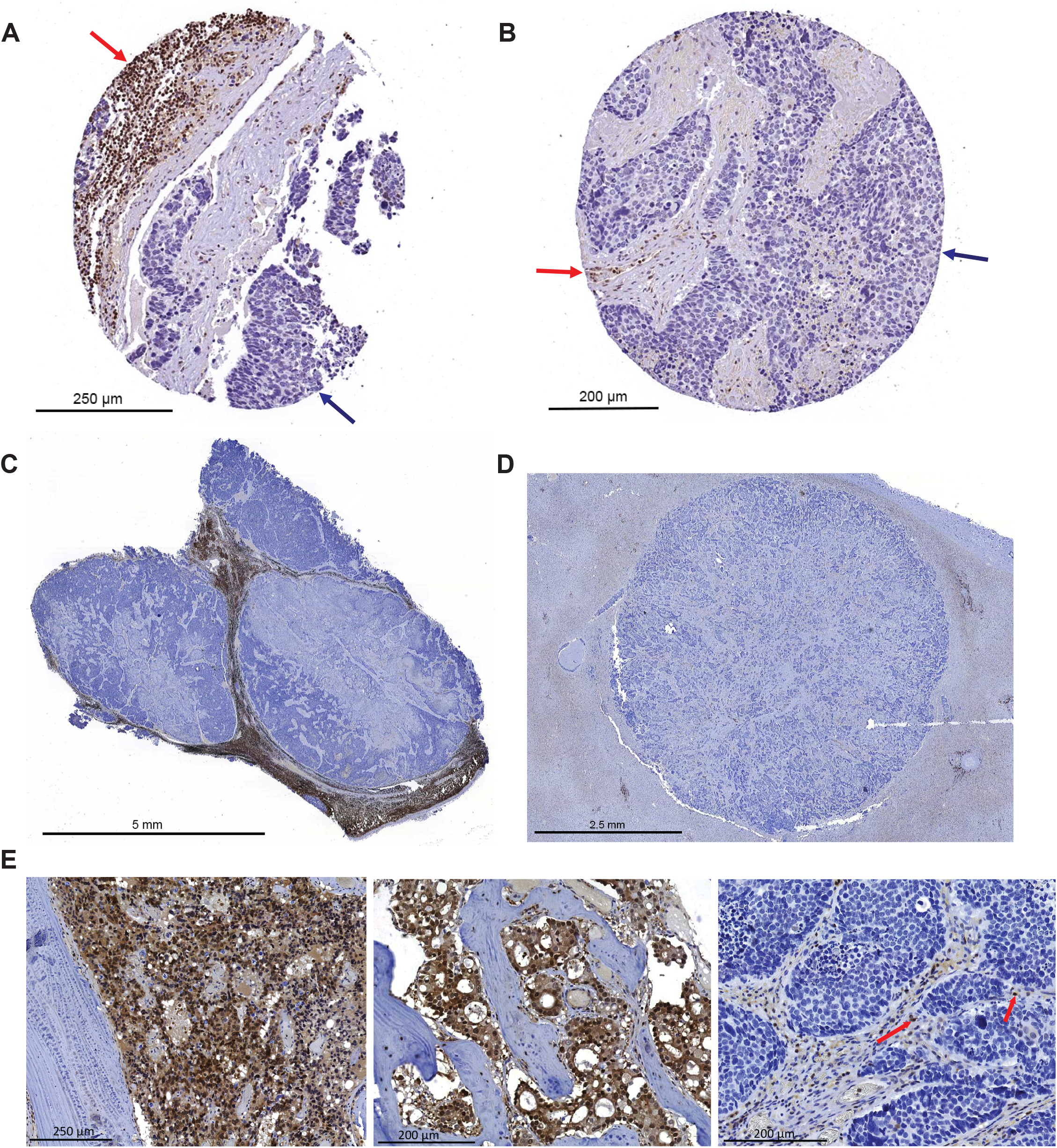
p27 IHC on metastatic prostate cancer TMA of 6 autopsy cases from 2 AA and 4 EA individuals. **A-B**) Examples of TMA spots exhibiting p27 loss in metastatic cancer cells in lymph node (A) and bone (B). p27 loss in metastatic cancer cells (blue arrows) and p27-intact lymphocytes (red arrows), indicate lack of p27 staining in metastatic cancer cells is not due to poor tissue fixation, warm ischemia time, or bone decalcification. **C-D**) p27 IHC on whole tissue sections from metastatic prostate cancer in a lymph node (C) and in liver (D). p27 loss in the metastatic niche is present in all cancer cells in the metastatic deposit. **E**) p27 IHC on whole tissue sections from metastatic prostate cancer in bone. The first two examples are p27+ metastatic cancer and the third example (far right) is a p27-bone metastasis. Red arrows denote p27-positive normal cells indicating that the lack of staining in p27-negative metastatic cancer cells is not due to poor tissue fixation or bone decalcification.

## Discussion

Collectively, our studies herein reveal that IPPL, that was tightly linked to biallelic genomic alterations by somatic mutation and/or deletion in *CDKN1B*, is an unexpectedly common occurrence in primary prostate cancer, with subclonal complete loss observed in close to 1 out of every 5 cancers that we assessed in AA individuals. This finding would place p27 as one of the most frequently mutated/deleted tumor suppressor genes known to occur in primary prostate cancer in AA males. Furthermore, whereas common somatic genomic alterations within tumor suppressor genes that are associated with worse outcomes in prostate cancer (TP53 mutation, PTEN loss) are often less common in primary prostate cancer in AA males, we report that subclonal *CDKN1B* loss is associated with worse outcomes and is also more prevalent in prostate cancer in AA males versus EA males. Of note, it is possible that our strategy of examining whole tissue sections may still underestimate the true prevalence of IPPL, as we only screened one slide containing the representative index tumor, and did not examine every tumor-containing block for each case. Future studies investigating p27 loss across the entirety of the cancer regions for each case are warranted.

One key finding from our study is that IPPL was almost always present as a presumably subclonal event in primary tumors. Panel sequencing of microdissected p27+ versus p27-cancer regions (**Fig. 2A, Table S1**) indicated stepwise mutation/loss of p27 alleles (with single copy loss in adjacent p27+ cancer and complete loss in p27-cancer) in multiple cases. Of the 63 cases identified with IPPL in our study, only two had larger p27-cancer regions than p27+ cancer regions, and both of those cases also had adjacent p27+ cancer, indicating a subclonal event (**Fig. S4**). Indeed, panel sequencing was performed on one of these cases (case 3, **Table S1**) and homozygous deletion of *CDKN1B* was detected in the p27-region, but not in the adjacent p27+ cancer.

In contrast, our analysis of metastatic prostate cancer indicated that all cells within a metastatic site can be p27− (**Fig 3C,D**). Likewise, the prevalence of p27 loss in metastatic disease is higher than that of primary prostate cancer in both publicly available datasets (**Table S2**) as well as our analysis of a limited number of autopsy cases (**Table S4**). These findings indicate that p27 loss may be a genomic alteration that is selected for in metastatic disease. We noted that most of the autopsy cases that we examined were heterogeneous in terms of exhibiting p27+ cancer at some metastatic sites, and p27-cancer in others. There was not a clear association between p27 loss in metastases and bone versus soft tissue sites of metastases (**Table S4**). We note that the reported frequency of mutations/deep deletions in *CDKN1B* in metastatic prostate cancer is far lower in cBioPortal datasets (average 7.6%, **Table S2**) than what we observed via p27 IHC in the autopsy TMA (4 of 6 cases assessed). We propose several possible reasons for this discrepancy. First, we screened the autopsy cases using p27 IHC. We confirmed that the protein loss that we observed in the metastatic cancer cells was not due to factors such as poor tissue fixation, warm ischemia time, or bone decalcification by examining non-cancer cells withing the metastatic niche and confirming expected nuclear p27 positivity in these cells (**Fig. 3**). We did not confirm genomic loss of *CDKN1B* in these cases however, and it is possible that p27 loss in metastatic disease is regulated by additional mechanisms such as epigenetic silencing or post-transcriptional regulation. This will be the focus of future studies. Another potential reason for the underestimation of *CDKN1B* biallelic alterations in metastases in cBioPortal datasets is related to our finding of heterogeneity in p27+ versus p27-metastatic tumors across the multiple metastatic sites sampled during autopsy (**Table S4**). The studies that provided the datasets in cBioPortal are typically restricted to one sample per patient, and therefore the chances of sampling a p27-metastatic site would be lower. Finally, tumor purity and the presence of normal cells can influence the detection of p27 genomic alterations. Likewise, cBioPortal reports deep deletions in *CDKN1B*, however our analysis in primary tumors indicated that *CDKN1B* alterations are often presumably hemizygous (**Fig. 2**).

There are some key limitations to our studies. Our strategy for laser microdissection of p27-regions increases tumor purity, however it does not eliminate normal cells such as stromal cells that have presumably genomically intact p27. As such, our ability to confidently distinguish heterozygous versus homozygous loss is limited in our panel sequencing analyses. Our interpretation of heterozygous loss in the data represented in **Fig. 2A** was therefore also guided by the status of p27 protein expression via IHC (**Table S1**). Although we screened a large number of primary prostatectomy cases for p27 subclonal loss using IHC (n=412) the number of outcome events for BCR and metastases were relatively low in this cohort and limited our risk assessments, especially in multivariate analyses. Future studies in larger cohorts linked to outcomes are warranted. Finally, all of our analyses were based on self-reported race, and although our prior studies in cohorts at our institution find a high association between estimated percentage of Yoruba in Ibadan, Nigeria (YRI) ancestry and self-reported AA race^47^, we did not assess specific associations between p27 loss and percent African ancestry in the present study.

We do not know why the association between IPPL and advanced pathologic stage as well as adverse outcomes (biochemical recurrence and metastases) was only significant among self-identified AA participants. We speculate that there may be other genomic or epigenomic alterations that co-occur with IPPL in these individuals that contribute to worse outcomes. Likewise, chronic inflammation is implicated in promoting tumor progression and metastasis, and prior work from our group and others suggest that prostate cancer arising in AA and EA males have differing inflammatory microenvironments, with higher expression of genes involved in inflammatory pathways in tumors from AA males than those of EA males.^13,47–54^ It is possible that chronic inflammation may cooperate with IPPL to promote metastases.

Overall, our study reveals that IPPL in primary prostate cancer is one of the most commonly occurring genomic alterations in known tumor suppressor genes to date, and one of the few known alterations that contribute to worse prostate cancer outcomes specifically in AA males.

## Acknowledgements

This work was supported by the Oncology Tissue and Imaging Servies core facility and the Molecular Diagnostics Laboratory in the Department of Pathology at Johns Hopkins Hospital. We would like to acknowledge Dr. Bruce Trock for assistance with outcomes data and Dr. Christopher Gocke for assistance with panel sequencing. We gratefully acknowledge grant support from Prostate Cancer Foundation Challenge Award 19CHAS03, Department of Defense, Prostate Cancer Research Program (PCRP) Award W81XWH-17-1-0286, and NIH R01 1R01CA288760.

## Figure Legends

**Figure S1**. Example of p27− cancer detected via IHC. Strong staining in p27+ normal cells in the stroma such as lymphocytes (arrows) indicates that the lack of signal in cancer cells is not due to poor tissue fixation.

**Figure S2**. Examples of mis-localized p27 expression as assessed by p27 IHC in prostate cancer. **A**) Normal prostate epithelium exhibiting normal nuclear p27 protein localization. **B**) Prostate cancer with predominantly nuclear p27 localization. **C**) Examples of lowered and predominantly cytoplasmic localization of p27 in p27+ cancer areas of case 1, 2, and 7 that contain presumably hemizygous genomic alterations to the *CDKN1B* gene. **D**) Example of lowered and predominantly cytoplasmic localization of p27 in p27+ cancer area of case 6 with no detected genomic alterations to the *CDKN1B* gene. Strong staining in p27+ normal cells in the stroma such as lymphocytes and endothelial cells (arrows) indicates that the lowered/mislocalized signal in cancer cells is not due to poor tissue fixation.

**Figure S3**. Example of p27− region (blue outline) that was missed during sampling for construction of a tissue microarray (TMA). TMA cores sampling only p27+ regions are denoted by arrows.

**Figure S4**. Examples of cases with expansive areas of p27 loss (blue outlined regions) that still have adjacent/admixed p27+ cancer regions (black outlined regions).

## References

1. Powell IJ, Bollig-Fischer A. Minireview: the molecular and genomic basis for prostate cancer health disparities. Mol Endocrinol. 2013;27:879–891.

2. DeSantis CE, Siegel RL, Sauer AG, Miller KD, Fedewa SA, Alcaraz KI, Jemal A. Cancer statistics for African Americans, 2016: Progress and opportunities in reducing racial disparities. CA Cancer J Clin. 2016;66:290–308.

3. Huang FW, Mosquera JM, Garofalo A, Oh C, Baco M, Amin-Mansour A, Rabasha B, Bahl S, Mullane SA, Robinson BD, Aldubayan S, Khani F, Karir B, Kim E, Chimene-Weiss J, Hofree M, Romanel A, Osborne JR, Kim JW, Azabdaftari G, Woloszynska-Read A, Sfanos K, De Marzo AM, Demichelis F, Gabriel S, Van Allen EM, Mesirov J, Tamayo P, Rubin MA, Powell IJ, Garraway LA. Exome sequencing of African-American prostate cancer reveals loss-of-function ERF mutations. Cancer Discov. 2017;7:973–983.

4. Koga Y, Song H, Chalmers ZR, Newberg J, Kim E, Carrot-Zhang J, Piou D, Polak P, Abdulkadir SA, Ziv E, Meyerson M, Frampton GM, Campbell JD, Huang FW. Genomic profiling of prostate cancers from men with African and European ancestry. Clin Cancer Res. 2020;26:4651–4660.

5. Mahal BA, Alshalalfa M, Kensler KH, Chowdhury-Paulino I, Kantoff P, Mucci LA, Schaeffer EM, Spratt D, Yamoah K, Nguyen PL, Rebbeck TR. Racial differences in genomic profiling of prostate cancer. N Engl J Med. 2020;383:1083–1085.

6. Pettersson A, Graff RE, Bauer SR, Pitt MJ, Lis RT, Stack EC, Martin NE, Kunz L, Penney KL, Ligon AH, Suppan C, Flavin R, Sesso HD, Rider JR, Sweeney C, Stampfer MJ, Fiorentino M, Kantoff PW, Sanda MG, Giovannucci EL, Ding EL, Loda M, Mucci LA. The TMPRSS2:ERG rearrangement, ERG expression, and prostate cancer outcomes: A cohort study and meta-analysis. Cancer Epidemiol Biomarkers Prev. 2012;21:1497–1509.

7. Magi-Galluzzi C, Tsusuki T, Elson P, Simmerman K, LaFargue C, Esgueva R, Klein E, Rubin MA, Zhou M. TMPRSS2-ERG gene fusion prevalence and class are significantly different in prostate cancer of Caucasian, African-American and Japanese patients. Prostate. 2011;71:489–497.

8. Khani F, Mosquera JM, Park K, Blattner M, O’Reilly C, MacDonald TY, Chen Z, Srivastava A, Tewari AK, Barbieri CE, Rubin MA, Robinson BD. Evidence for molecular differences in prostate cancer between African American and Caucasian men. Clin Cancer Res. 2014;20:4925–4934.

9. Stopsack KH, Nandakumar S, Arora K, Nguyen B, Vasselman SE, Nweji B, McBride SM, Morris MJ, Rathkopf DE, Slovin SF, Danila DC, Autio KA, Scher HI, Mucci LA, Solit DB, Gönen M, Chen Y, Berger MF, Schultz N, Abida W, Kantoff PW. Differences in prostate cancer genomes by self-reported race: Contributions of genetic ancestry, modifiable cancer risk factors, and clinical factors. Clin. Cancer Res. 2022;28:318–326.

10. Tosoian JJ, Almutairi F, Morais CL, Glavaris S, Hicks J, Sundi D, Humphreys E, Han M, De Marzo AM, Ross AE, Tomlins SA, Schaeffer EM, Trock BJ, Lotan TL. Prevalence and prognostic significance of PTEN loss in African-American and European-American men undergoing radical prostatectomy. Eur Urol. 2017;71:697–700.

11. Liu W, Zheng SL, Na R, Wei L, Sun J, Gallagher J, Wei J, Resurreccion WK, Ernst S, Sfanos KS, Isaacs WB, Xu J. Distinct genomic alterations in prostate tumors derived from African American men. Mol Cancer Res. 2020;18:1815–1824.

12. Yuan J, Kensler KH, Hu Z, Zhang Y, Zhang T, Jiang J, Xu M, Pan Y, Long M, Montone KT, Tanyi JL, Fan Y, Zhang R, Hu X, Rebbeck TR, Zhang L. Integrative comparison of the genomic and transcriptomic landscape between prostate cancer patients of predominantly African or European genetic ancestry. PLoS Genet. 2020;16:e1008641.

13. Vidal I, Zheng Q, Hicks JL, Chen J, Platz EA, Trock BJ, Kulac I, Baena-Del Valle JA, Sfanos KS, Ernst S, Jones T, Maynard JP, Glavaris SA, Nelson WG, Yegnasubramanian S, De Marzo AM. GSTP1 positive prostatic adenocarcinomas are more common in Black than White men in the United States. PLoS One. 2021;16:e0241934.

14. Petrovics G, Li H, Stümpel T, Tan S-H, Young D, Katta S, Li Q, Ying K, Klocke B, Ravindranath L, Kohaar I, Chen Y, Ribli D, Grote K, Zou H, Cheng J, Dalgard CL, Zhang S, Csabai I, Kagan J, Takeda D, Loda M, Srivastava S, Scherf M, Seifert M, Gaiser T, McLeod DG, Szallasi Z, Ebner R, Werner T, Sesterhenn IA, Freedman M, Dobi A, Srivastava S. A novel genomic alteration of LSAMP associates with aggressive prostate cancer in African American men. EBioMedicine. 2015;2:1957–1964.

15. Sharma SS, Pledger WJ. The non-canonical functions of p27(Kip1) in normal and tumor biology. Cell Cycle. 2016;15:1189–1201.

16. Fero ML, Randel E, Gurley KE, Roberts JM, Kemp CJ. The murine gene p27Kip1 is haplo-insufficient for tumour suppression. Nature. 1998;396:177–180.

17. Stephens PJ, Tarpey PS, Davies H, Van Loo P, Greenman C, Wedge DC, Nik-Zainal S, Martin S, Varela I, Bignell GR, Yates LR, Papaemmanuil E, Beare D, Butler A, Cheverton A, Gamble J, Hinton J, Jia M, Jayakumar A, Jones D, Latimer C, Lau KW, McLaren S, McBride DJ, Menzies A, Mudie L, Raine K, Rad R, Spencer Chapman M, Teague J, Easton D, Langerød A, Lee MTM, Shen C-Y, Tee BTK, Huimin BW, Broeks A, Vargas AC, Turashvili G, Martens J, Fatima A, Miron P, Chin S-F, Thomas G, Boyault S, Mariani O, Lakhani SR, van de Vijver M, van ‘t Veer L, Foekens J, Desmedt C, Sotiriou C, Tutt A, Caldas C, Reis-Filho JS, Aparicio SAJR, Salomon AV, Børresen-Dale A-L, Richardson AL, Campbell PJ, Futreal PA, Stratton MR. The landscape of cancer genes and mutational processes in breast cancer. Nature. 2012;486:400–404.

18. Viotto D, Russo F, Anania I, Segatto I, Rampioni Vinciguerra GL, Dall’Acqua A, Bomben R, Perin T, Cusan M, Schiappacassi M, Gerratana L, D’Andrea S, Citron F, Vit F, Musco L, Mattevi MC, Mungo G, Nicoloso MS, Sonego M, Massarut S, Sorio R, Barzan L, Franchin G, Giorda G, Lucia E, Sulfaro S, Giacomarra V, Polesel J, Toffolutti F, Canzonieri V, Puglisi F, Gattei V, Vecchione A, Belletti B, Baldassarre G. CDKN1B mutation and copy number variation are associated with tumor aggressiveness in luminal breast cancer. J Pathol. 2021;253:234–245.

19. Kossatz U, Malek NP. p27: tumor suppressor and oncogene …? Cell Res. 2007;17:832–833.

20. Tsihlias J, Kapusta L, Slingerland J. The prognostic significance of altered cyclin-dependent kinase inhibitors in human cancer. Annu Rev Med. 1999;50:401–423.

21. Besson A, Hwang HC, Cicero S, Donovan SL, Gurian-West M, Johnson D, Clurman BE, Dyer MA, Roberts JM. Discovery of an oncogenic activity in p27Kip1 that causes stem cell expansion and a multiple tumor phenotype. Genes Dev. 2007;21:1731–1746.

22. Wu FY, Wang SE, Sanders ME, Shin I, Rojo F, Baselga J, Arteaga CL. Reduction of cytosolic p27(Kip1) inhibits cancer cell motility, survival, and tumorigenicity. Cancer Res. 2006;66:2162–2172.

23. De Marzo AM, Meeker AK, Epstein JI, Coffey DS. Prostate stem cell compartments: expression of the cell cycle inhibitor p27Kip1 in normal, hyperplastic, and neoplastic cells. Am J Pathol. 1998;153:911–919.

24. Guo Y, Sklar GN, Borkowski A, Kyprianou N. Loss of the cyclin-dependent kinase inhibitor p27(Kip1) protein in human prostate cancer correlates with tumor grade. Clin Cancer Res. 1997;3:2269–2274.

25. Tsihlias J, Kapusta LR, DeBoer G, Morava-Protzner I, Zbieranowski I, Bhattacharya N, Catzavelos GC, Klotz LH, Slingerland JM. Loss of cyclin-dependent kinase inhibitor p27Kip1 is a novel prognostic factor in localized human prostate adenocarcinoma. Cancer Res. 1998;58:542–548.

26. Kluth M, Ahrary R, Hube-Magg C, Ahmed M, Volta H, Schwemin C, Steurer S, Wittmer C, Wilczak W, Burandt E, Krech T, Adam M, Michl U, Heinzer H, Salomon G, Graefen M, Koop C, Minner S, Simon R, Sauter G, Schlomm T. Genomic deletion of chromosome 12p is an independent prognostic marker in prostate cancer. Oncotarget. 2015;6:27966–27979.

27. Ananthanarayanan V, Deaton RJ, Amatya A, Macias V, Luther E, Kajdacsy-Balla A, Gann PH. Subcellular localization of p27 and prostate cancer recurrence: automated digital microscopy analysis of tissue microarrays. Hum Pathol. 2011;42:873–881.

28. Cordon-Cardo C, Koff A, Drobnjak M, Capodieci P, Osman I, Millard SS, Gaudin PB, Fazzari M, Zhang ZF, Massague J, Scher HI. Distinct altered patterns of p27KIP1 gene expression in benign prostatic hyperplasia and prostatic carcinoma. J Natl Cancer Inst. 1998;90:1284–1291.

29. Yang RM, Naitoh J, Murphy M, Wang HJ, Phillipson J, deKernion JB, Loda M, Reiter RE. Low p27 expression predicts poor disease-free survival in patients with prostate cancer. J Urol. 1998;159:941–945.

30. Kuczyk M, Machtens S, Hradil K, Schubach J, Christian W, Knüchel R, Hartmann J, Bokemeyer C, Jonas U, Serth J. Predictive value of decreased p27Kip1 protein expression for the recurrence-free and long-term survival of prostate cancer patients. Br J Cancer. 1999;81:1052–1058.

31. Di Cristofano A, De Acetis M, Koff A, Cordon-Cardo C, Pandolfi PP. Pten and p27KIP1 cooperate in prostate cancer tumor suppression in the mouse. Nat Genet. 2001;27:222–224.

32. Halvorsen OJ, Haukaas SA, Akslen LA. Combined loss of PTEN and p27 expression is associated with tumor cell proliferation by Ki-67 and increased risk of recurrent disease in localized prostate cancer. Clin Cancer Res. 2003;9:1474–1479.

33. Kibel AS, Schutte M, Kern SE, Isaacs WB, Bova GS. Identification of 12p as a region of frequent deletion in advanced prostate cancer. Cancer Res. 1998;58:5652–5655.

34. Quigley DA, Dang HX, Zhao SG, Lloyd P, Aggarwal R, Alumkal JJ, Foye A, Kothari V, Perry MD, Bailey AM, Playdle D, Barnard TJ, Zhang L, Zhang J, Youngren JF, Cieslik MP, Parolia A, Beer TM, Thomas G, Chi KN, Gleave M, Lack NA, Zoubeidi A, Reiter RE, Rettig MB, Witte O, Ryan CJ, Fong L, Kim W, Friedlander T, Chou J, Li H, Das R, Li H, Moussavi-Baygi R, Goodarzi H, Gilbert LA, Lara PN Jr, Evans CP, Goldstein TC, Stuart JM, Tomlins SA, Spratt DE, Cheetham RK, Cheng DT, Farh K, Gehring JS, Hakenberg J, Liao A, Febbo PG, Shon J, Sickler B, Batzoglou S, Knudsen KE, He HH, Huang J, Wyatt AW, Dehm SM, Ashworth A, Chinnaiyan AM, Maher CA, Small EJ, Feng FY. Genomic hallmarks and structural variation in metastatic prostate cancer. Cell. 2018;174:758-769.e9.

35. Annala M, Fu S, Bacon JVW, Sipola J, Iqbal N, Ferrario C, Ong M, Wadhwa D, Hotte SJ, Lo G, Tran B, Wood LA, Gingerich JR, North SA, Pezaro CJ, Ruether JD, Sridhar SS, Kallio HML, Khalaf DJ, Wong A, Beja K, Schönlau E, Taavitsainen S, Nykter M, Vandekerkhove G, Azad AA, Wyatt AW, Chi KN. Cabazitaxel versus abiraterone or enzalutamide in poor prognosis metastatic castration-resistant prostate cancer: a multicentre, randomised, open-label, phase II trial. Ann Oncol. 2021;32:896–905.

36. Wyatt AW, Annala M, Aggarwal R, Beja K, Feng F, Youngren J, Foye A, Lloyd P, Nykter M, Beer TM, Alumkal JJ, Thomas GV, Reiter RE, Rettig MB, Evans CP, Gao AC, Chi KN, Small EJ, Gleave ME. Concordance of circulating tumor DNA and matched metastatic tissue biopsy in prostate cancer. J Natl Cancer Inst [Internet]. 2017;109. Available from: 10.1093/jnci/djx118

37. Faisal FA, Murali S, Kaur H, Vidotto T, Guedes LB, Salles DC, Kothari V, Tosoian JJ, Han S, Hovelson DH, Hu K, Spratt DE, Baras AS, Tomlins SA, Schaeffer EM, Lotan TL. CDKN1B deletions are associated with metastasis in African American men with clinically localized, surgically treated prostate cancer. Clin Cancer Res. 2020;26:2595–2602.

38. Abida W, Armenia J, Gopalan A, Brennan R, Walsh M, Barron D, Danila D, Rathkopf D, Morris M, Slovin S, McLaughlin B, Curtis K, Hyman DM, Durack JC, Solomon SB, Arcila ME, Zehir A, Syed A, Gao J, Chakravarty D, Vargas HA, Robson ME, Joseph V, Offit K, Donoghue MTA, Abeshouse AA, Kundra R, Heins ZJ, Penson AV, Harris C, Taylor BS, Ladanyi M, Mandelker D, Zhang L, Reuter VE, Kantoff PW, Solit DB, Berger MF, Sawyers CL, Schultz N, Scher HI. Prospective genomic profiling of prostate cancer across disease states reveals germline and somatic alterations that may affect clinical decision making. JCO Precis Oncol [Internet]. 2017;2017. Available from: 10.1200/PO.17.00029

39. Robinson D, Van Allen EM, Wu Y-M, Schultz N, Lonigro RJ, Mosquera J-M, Montgomery B, Taplin M-E, Pritchard CC, Attard G, Beltran H, Abida W, Bradley RK, Vinson J, Cao X, Vats P, Kunju LP, Hussain M, Feng FY, Tomlins SA, Cooney KA, Smith DC, Brennan C, Siddiqui J, Mehra R, Chen Y, Rathkopf DE, Morris MJ, Solomon SB, Durack JC, Reuter VE, Gopalan A, Gao J, Loda M, Lis RT, Bowden M, Balk SP, Gaviola G, Sougnez C, Gupta M, Yu EY, Mostaghel EA, Cheng HH, Mulcahy H, True LD, Plymate SR, Dvinge H, Ferraldeschi R, Flohr P, Miranda S, Zafeiriou Z, Tunariu N, Mateo J, Perez-Lopez R, Demichelis F, Robinson BD, Schiffman M, Nanus DM, Tagawa ST, Sigaras A, Eng KW, Elemento O, Sboner A, Heath EI, Scher HI, Pienta KJ, Kantoff P, de Bono JS, Rubin MA, Nelson PS, Garraway LA, Sawyers CL, Chinnaiyan AM. Integrative clinical genomics of advanced prostate cancer. Cell. 2015;161:1215–1228.

40. Armenia J, Wankowicz SAM, Liu D, Gao J, Kundra R, Reznik E, Chatila WK, Chakravarty D, Han GC, Coleman I, Montgomery B, Pritchard C, Morrissey C, Barbieri CE, Beltran H, Sboner A, Zafeiriou Z, Miranda S, Bielski CM, Penson AV, Tolonen C, Huang FW, Robinson D, Wu YM, Lonigro R, Garraway LA, Demichelis F, Kantoff PW, Taplin M-E, Abida W, Taylor BS, Scher HI, Nelson PS, de Bono JS, Rubin MA, Sawyers CL, Chinnaiyan AM, PCF/SU2C International Prostate Cancer Dream Team, Schultz N, Van Allen EM. The long tail of oncogenic drivers in prostate cancer. Nat Genet. 2018;50:645– 651.

41. Taylor BS, Schultz N, Hieronymus H, Gopalan A, Xiao Y, Carver BS, Arora VK, Kaushik P, Cerami E, Reva B, Antipin Y, Mitsiades N, Landers T, Dolgalev I, Major JE, Wilson M, Socci ND, Lash AE, Heguy A, Eastham JA, Scher HI, Reuter VE, Scardino PT, Sander C, Sawyers CL, Gerald WL. Integrative genomic profiling of human prostate cancer. Cancer Cell. 2010;18:11–22.

42. Maynard JP, Lu J, Vidal I, Hicks J, Mummert L, Ali T, Kempski R, Carter AM, Sosa RY, Peiffer LB, Joshu CE, Lotan TL, De Marzo AM, Sfanos KS. P2X4 purinergic receptors offer a therapeutic target for aggressive prostate cancer. J Pathol. 2022;256:149–163.

43. De Marzo AM, Fedor HH, Gage WR, Rubin MA. Inadequate formalin fixation decreases reliability of p27 immunohistochemical staining: probing optimal fixation time using high-density tissue microarrays. Hum Pathol. 2002;33:756–760.

44. Kaur HB, Lu J, Guedes LB, Maldonado L, Reitz L, Barber JR, De Marzo AM, Tomlins SA, Sfanos KS, Eisenberger M, Schaeffer EM, Joshu CE, Lotan TL. TP53 missense mutation is associated with increased tumor-infiltrating T cells in primary prostate cancer. Hum Pathol. 2019;87:95–102.

45. Kaur HB, Guedes LB, Lu J, Maldonado L, Reitz L, Barber JR, De Marzo AM, Tosoian JJ, Tomlins SA, Schaeffer EM, Joshu CE, Sfanos KS, Lotan TL. Association of tumor-infiltrating T-cell density with molecular subtype, racial ancestry and clinical outcomes in prostate cancer. Mod Pathol. 2018;31:1539–1552.

46. Cerami E, Gao J, Dogrusoz U, Gross BE, Sumer SO, Aksoy BA, Jacobsen A, Byrne CJ, Heuer ML, Larsson E, Antipin Y, Reva B, Goldberg AP, Sander C, Schultz N. The cBio cancer genomics portal: an open platform for exploring multidimensional cancer genomics data. Cancer Discov. 2012;2:401–404.

47. Mendes AA, Lu J, Kaur HB, Zheng SL, Xu J, Hicks J, Weiner AB, Schaeffer EM, Ross AE, Balk SP, Taplin M-E, Lack NA, Tekoglu E, Maynard JP, De Marzo AM, Antonarakis ES, Sfanos KS, Joshu CE, Shenderov E, Lotan TL. Association of B7-H3 expression with racial ancestry, immune cell density, and androgen receptor activation in prostate cancer. Cancer. 2022;128:2269–2280.

48. Hempel Sullivan H, Heaphy CM, Kulac I, Cuka N, Lu J, Barber JR, De Marzo AM, Lotan TL, Joshu CE, Sfanos KS. High extratumoral mast cell counts are associated with a higher risk of adverse prostate cancer outcomes. Cancer Epidemiol Biomarkers Prev. 2020;29:668–675.

49. Maynard JP, Godwin TN, Lu J, Vidal I, Lotan TL, De Marzo AM, Joshu CE, Sfanos KS. Localization of macrophage subtypes and neutrophils in the prostate tumor microenvironment and their association with prostate cancer racial disparities. Prostate. 2022;82:1505–1519.

50. Eastham JA, May RA, Whatley T, Crow A, Venable DD, Sartor O. Clinical characteristics and biopsy specimen features in African-American and white men without prostate cancer. J Natl Cancer Inst. 1998;90:756–760.

51. Powell IJ, Dyson G, Land S, Ruterbusch J, Bock CH, Lenk S, Herawi M, Everson R, Giroux CN, Schwartz AG, Bollig-Fischer A. Genes associated with prostate cancer are differentially expressed in African American and European American men. Cancer Epidemiol Biomarkers Prev. 2013;22:891–897.

52. Reams RR, Agrawal D, Davis MB, Yoder S, Odedina FT, Kumar N, Higginbotham JM, Akinremi T, Suther S, Soliman KF. Microarray comparison of prostate tumor gene expression in African-American and Caucasian American males: a pilot project study. Infect Agent Cancer. 2009;4 Suppl 1:S3.

53. Wallace TA, Prueitt RL, Yi M, Howe TM, Gillespie JW, Yfantis HG, Stephens RM, Caporaso NE, Loffredo CA, Ambs S. Tumor immunobiological differences in prostate cancer between African-American and European-American men. Cancer Res. 2008;68:927– 936.

54. Hardiman G, Savage SJ, Hazard ES, Wilson RC, Courtney SM, Smith MT, Hollis BW, Halbert CH, Gattoni-Celli S. Systems analysis of the prostate transcriptome in African-American men compared with European-American men. Pharmacogenomics. 2016;17:1129–1143.

