## Supplementary figures and images for "Subclonal Complete Loss of *CDKN1B* as a Common Genomic Alteration in Prostate Cancer: Associations with Race and Prostate Cancer Outcomes"

### Supplemental Figures S1-S4

#### Figure S1

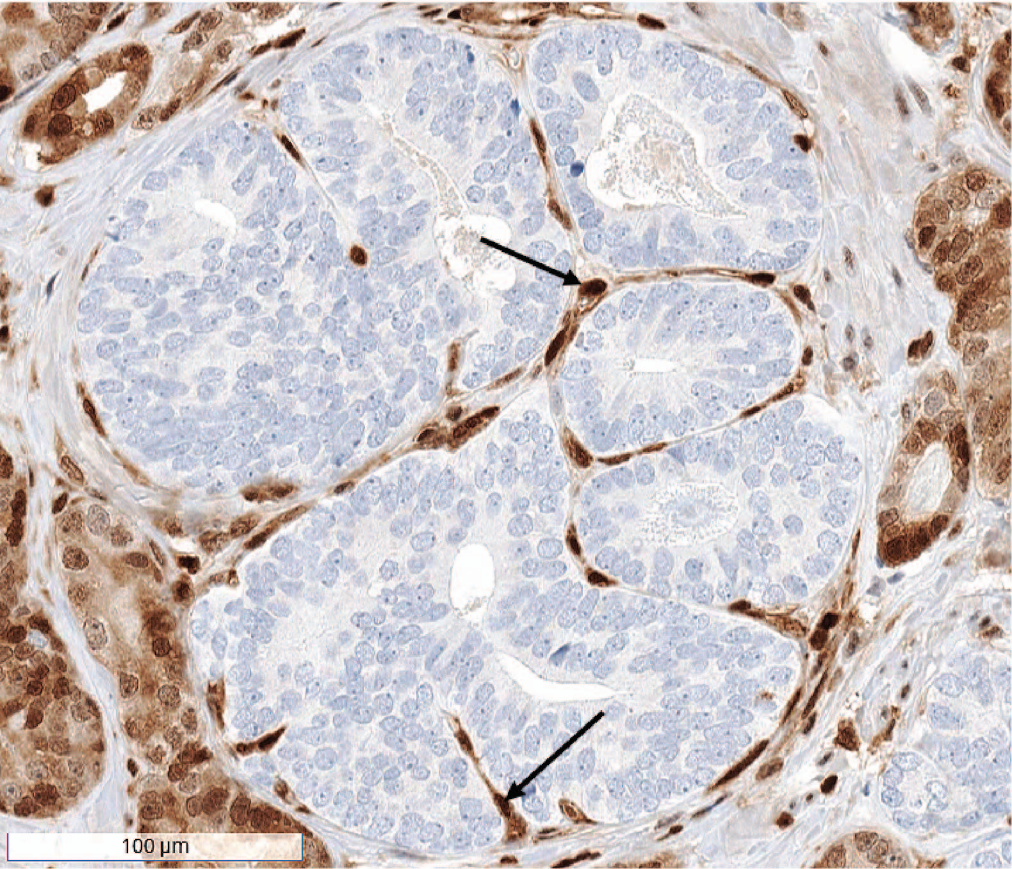

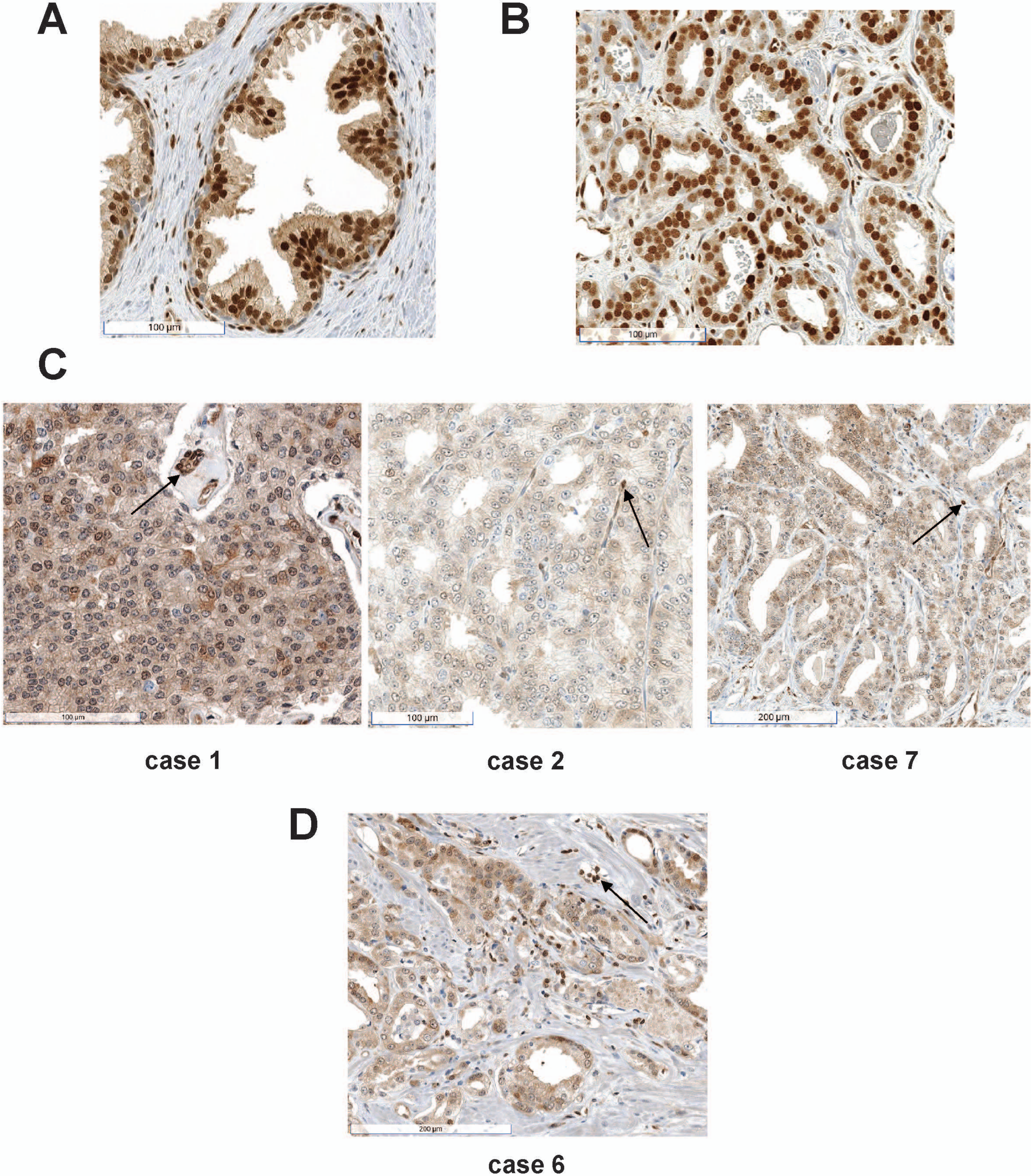

#### Figure S3

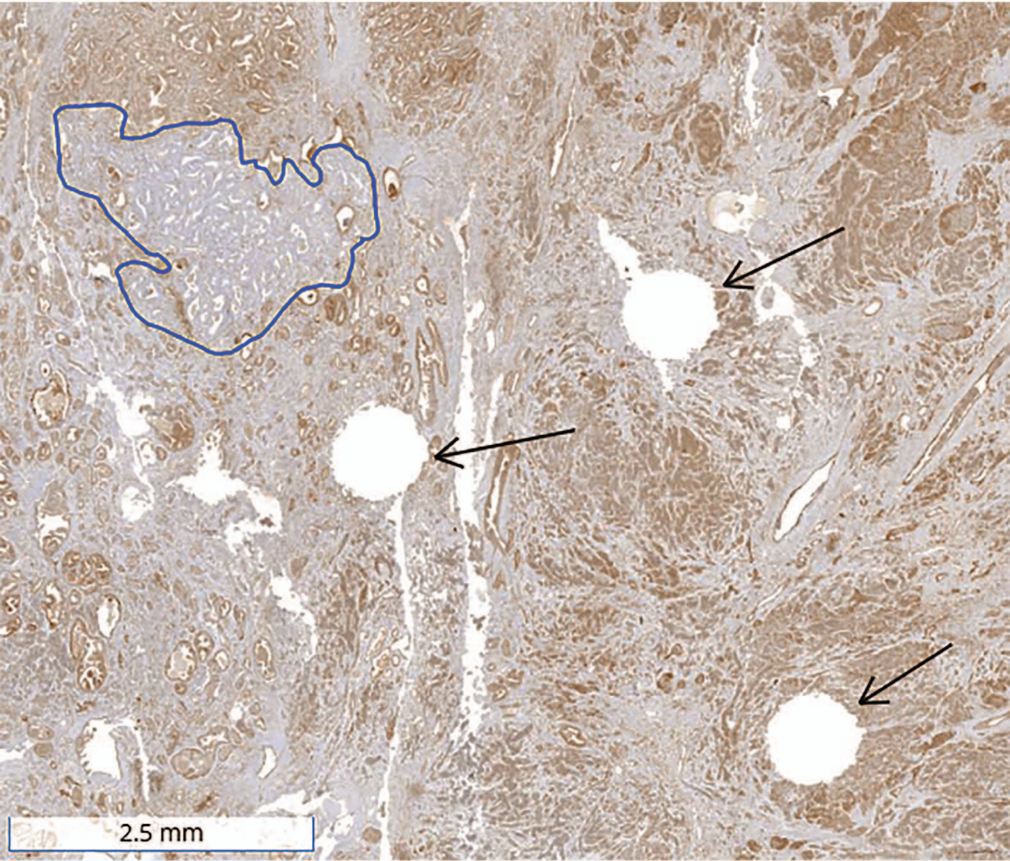

**Figure S4**

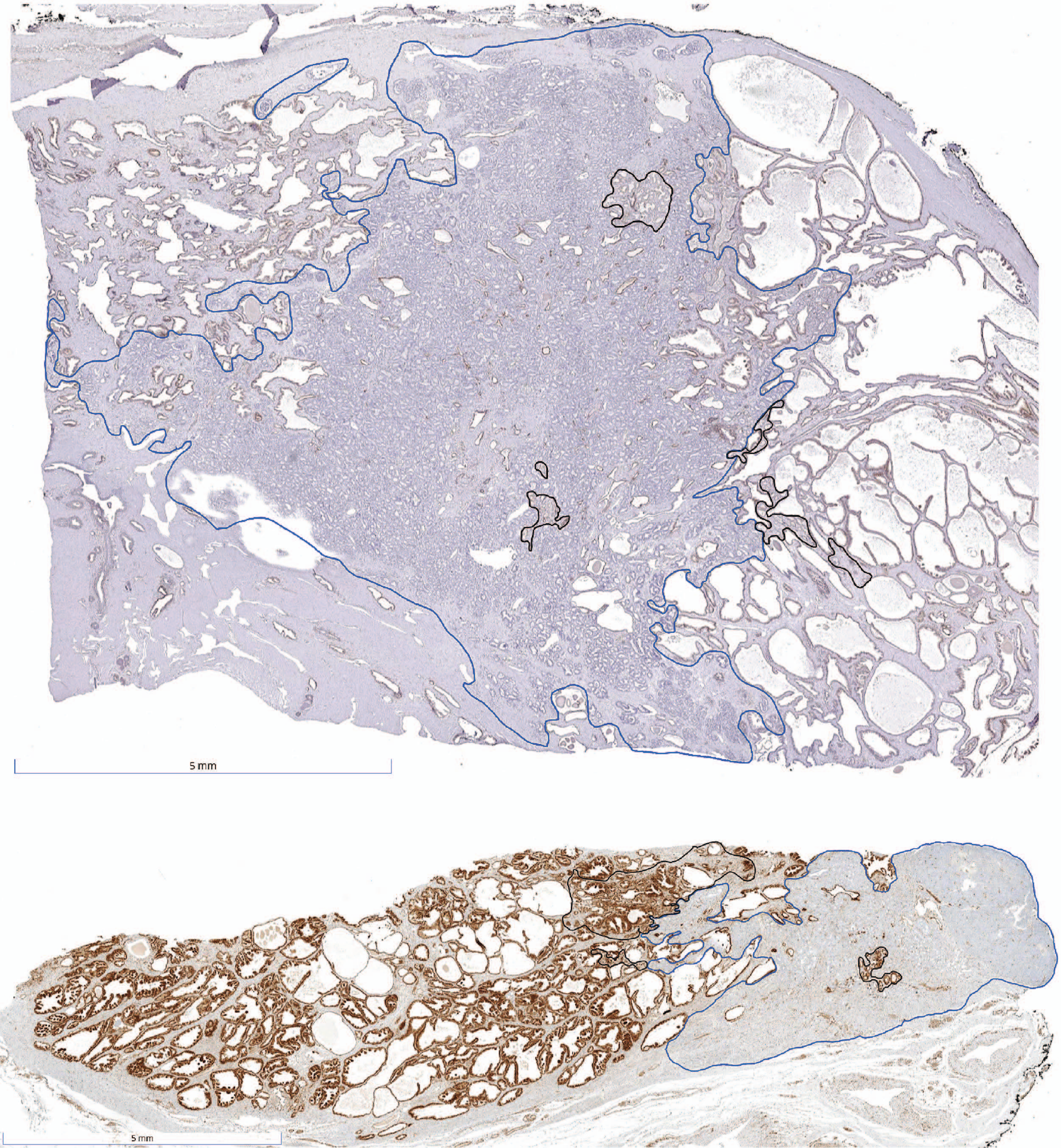
