## Supplemental Table 5 for "Subclonal Complete Loss of *CDKN1B* as a Common Genomic Alteration in Prostate Cancer: Associations with Race and Prostate Cancer Outcomes"

**Supplemental Table S5**. Comparison of p27 loss status to other common genomic alterations in primary prostate cancer.

| | All | | | | | AA | | | | | EA | | | | |
|---|---|---|---|---|---|---|---|---|---|---|---|---|---|---|---|
| | Number of Patients | | | | p * | Number of Patients | | | | p * | Number of Patients | | | | p * |
| | P27 Intact | | P27 Loss | | | P27 Intact | | P27 Loss | | | P27 Intact | | P27 Loss | | |
| PTEN | | | | | | | | | | | | | | | |
| Intact | 193 | (82.13) | 32 | (82.05) | 0.9 | 119 | (85.00) | 26 | (83.87) | 0.8 | 73 | (77.66) | 6 | (75.00) | 0.9 |
| Loss | 42 | (17.87) | 7 | (17.95) | | 21 | (15.00) | 5 | (16.13) | | 21 | (22.34) | 2 | (25.00) | |
| ERG | | | | | | | | | | | | | | | |
| Negative | 154 | (65.53) | 30 | (76.92) | 0.2 | 108 | (77.14) | 25 | (80.65) | 0.7 | 45 | (47.87) | 5 | (62.50) | 0.5 |
| Positive | 81 | (34.47) | 9 | (23.08) | | 32 | (22.86) | 6 | (19.35) | | 49 | (52.13) | 3 | (37.50) | |
| TP53 mutation | | | | | | | | | | | | | | | |
| Negative | 90 | (91.84) | 15 | (93.75) | 0.9 | 43 | (93.48) | 10 | (100.00) | 0.9 | 46 | (90.20) | 5 | (83.33) | 0.5 |
| Positive | 8 | (8.16) | 1 | (6.25) | | 3 | (6.52) | 0 | (0.00) | | 5 | (9.80) | 1 | (16.67) | |

* Chi-Square test or Fisher's Exact test
